# Grooming interventions in female rhesus macaques as social niche construction

**DOI:** 10.1101/2020.06.29.178004

**Authors:** Alexander Mielke, Carina Bruchmann, Oliver Schülke, Julia Ostner

**Affiliations:** Primate Models of Behavioural Evolution, Oxford University, Oxford, UK; Department for Behavioral Ecology, University of Goettingen, Goettingen, Germany; Primate Social Evolution Group, German Primate Center, Leibniz Institute for Primate Research, Goettingen, Germany; Leibniz ScienceCampus Primate Cognition, German Primate Center, Leibniz Institute for Primate Research, Goettingen, Germany

## Abstract

Social animals invest time and resources into building and adapting their social environment, which emerges not only from their own but also from the decisions of other group members. Thus, individuals have to monitor interactions between others and potentially decide when and how to interfere to prevent damage to their own investment. These interventions can be subtle, as in the case of affiliative interactions such as grooming, but they can inform us about how animals structure their world and influence other group members. Here, we used interventions into grooming bouts in 29 female rhesus macaques (*Macaca mulatta*) to determine who intervened into which grooming bouts, why, and what determined intervention outcomes, based on kinship, dominance rank, and affiliative relationships between groomers and (potential) interveners. Using 1132 grooming bouts and 521 interventions, we show that high dominance rank of groomers reduced the risk of intervention. Bystanders, particularly when high-ranking, intervened in grooming of their kin, close affiliates, and close-ranked competitors. Interveners gained access to their close affiliates for subsequent grooming. Affiliative relationship and rank determined intervention outcomes, with reduced aggression risk facilitating grooming involving three individuals. Thus, interventions in this species involved the monitoring of grooming interactions, decision-making based on several individual and dyadic characteristics, and potentially allowed individuals to broaden their access to grooming partners, protect their own relationships, and influence their social niche.

## Introduction and Hypotheses

Animals living in stable social groups must navigate a web of social relationships with group members daily. Some of these relationships, e.g. with kin and regular cooperation partners, are a central in the lives of many species, and forming and maintaining them has a measurable impact on individual fitness (Ostner & Schülke, 2018; Silk et al., 2013; Snyder-Mackler et al., 2020). Stable bonds seem to impact individual fitness by enhancing individual survival (Archie et al., 2014; Ellis et al., 2019), infant survival (Cameron et al., 2009; Silk et al., 2009), access to mating partners (Städele et al., 2019), food (Carter & Wilkinson, 2013; Samuni et al., 2018), and rank attainment (Schülke et al., 2010). Individuals will invest time and resources into forming bonds, which often remain stable across years (Kalbitz et al., 2016; Massen & Sterck, 2013; Silk et al., 2012). However, group social structure changes with demographic and dominance hierarchy changes, and individuals may compete over access to cooperation partners (Noë & Hammerstein, 1995).

This puts individuals in a bind: losing a bond partner means losing future support and past investment (Mielke et al., 2017). To secure their investment, individuals reconcile with their bonded partners after conflict (Cords & Aureli, 2000); often, however, it is not their own behaviour threatening the relationship, but actions of third parties. Therefore, we would predict that mechanisms evolve that allow individuals to protect their relationships against outside interference.

Interactions involving third parties in animals are remarkable as they show us that animals are active agents gathering and evaluating information about their social network and shaping the lives of those around them (Schülke et al., 2020; Seyfarth & Cheney, 2015). This process of individuals influencing interaction patterns around them has been termed ‘social nice construction’ (Barrett et al., 2012; Flack et al., 2006; Ryan et al., 2016): individuals’ behaviour changes the social landscape of the group and the selective context in which social behaviour occurs. By affecting how group members interact with each other, individuals influence their own future interactions (Ryan et al., 2016). Studies on third-party influence in social animals have focused on agonistic interactions and bystanders’ role in supporting fighters (Kajokaite et al., 2019; Young et al., 2014), policing fights (Beisner & McCowan, 2013; Flack et al., 2006), reconciling participants (Cords & Aureli, 2000), and consoling victims (de Waal & van Roosmalen, 1979; Preis et al., 2018). Monitoring interactions of other agents and considering the social environment when making decisions increases the information processing involved in successfully navigating a social group in everyday life, which may require a cognitive apparatus able to meet those challenges (Jolly, 1966).

In recent years, interest has moved towards socio-positive interactions, and the role bystanders play in changing partner choice (Mielke et al., 2018) and interaction outcome (Newton-Fisher & Kaburu, 2017). Bystanders can get involved by interfering in cooperation of others; previous studies have repeatedly shown that cooperation interventions disrupt bond formation if resulting changes in relationships would be detrimental to bystanders. In feral horses, individuals are more likely to disrupt affiliative interactions if one partner is a close affiliate (Schneider & Krueger, 2012; VanDierendonck et al., 2009). In dogs, play interventions target the ‘losing’ player independent of relationship status (Ward et al., 2009), and individuals show ‘jealous’ behaviour when their owner interacts with a dog model (Harris & Prouvost, 2014). Paired ravens intervene in pair formation attempts of new pairs, who will be more likely to compete for resources (Massen et al., 2014). Grooming interventions allow mandrills to restrict grooming access of lower-ranking competitors and limit alliance formation (Schino & Lasio, 2018). In stumptailed macaques, several affiliative interaction types see interventions (Mondragón-Ceballos, 2001). Sooty mangabeys and chimpanzees intervene in grooming to prevent their bond partners and close-ranked competitors from forming new connections (Mielke et al., 2017), and conversely they choose grooming partners to minimise the threat of interventions (Mielke et al., 2018).

We thus see a pattern emerging across species where third parties shape socio-positive interactions and the social niche an individual inhabits the same way they do in agonistic contexts. However, the number of species that we have information for is still limited, and the impact of many relevant factors unexplored. For example, the role of kin-relations in interventions is poorly understood: In theory, kin should not have to resort to interventions to protect affiliative relationships as these should be more stable than bonds among non-kin (Silk et al., 2010). Kin bonds should be sustained independent of investment. However, not all kin dyads cooperate at above-expected levels (De Moor et al., 2020), and in larger matrilines, kin could compete with each other for access to other related individuals, for example siblings fighting to gain access to their mother.

Here, we use rhesus macaques (*Macaca mulatta*) as a model to understand the decisions underlying grooming interventions. Female rhesus macaque societies are despotic, intolerant, and matrilineal structured (Thierry et al., 2008), but non-kin bonds of females exist and impact individual fitness (Ellis et al., 2019). Grooming and similar low-cost forms of cooperation (Carter et al., 2020) are the most visible behaviours used to negotiate bond maintenance and formation, and are a vital mechanism to negotiate social relationships in female rhesus macaques (Balasubramaniam & Berman, 2017). Grooming takes time and happens in the open, with group members watching, potentially giving them a way to influence cooperative exchanges. Females monitor who is close when interacting with offspring (Semple et al., 2009) and selectively attend to agonistic and affiliative interactions in their vicinity and particularly so if a close affiliate or a higher ranking female is involved (Schülke et al. 2020). By focusing on a group of females with known ranks, affiliative relationships, and kin relations, we can disentangle grooming interventions: a) Which dyads are most at risk of grooming interventions? b) If interventions occur, who intervenes? c) When intervening, whom of the groomers are interveners trying to get access to (the ‘target’ of the intervention)? d) What determines intervention outcomes? We predicted that, due to the despotic intolerant social structure of female rhesus macaques, rank plays an outsized role in deciding who can successfully intervene, but that the decision to intervene or not is driven by kinship and social bond strength between potential interveners and the groomers.

## Methods

### Study Group and Observations

For this study, C.B. observed the 29 adult females aged above 3 years in one group of 39 rhesus macaques with a single adult male at the German Primate Center, Goettingen between January and March 2019. All individuals were identifiable by a tattooed four-number code on their chest and individual differences in fur coloration, size, and natural markings. The group had free access to a 250sqm outdoor and a 48sqm indoor enclosure, were fed once a day with fresh fruit and vegetables, once with a cottage cheese and grain preparation and once with monkey chow and had ad libitum access to water from several faucets.

Data were collected using three different methods. A 30min instantaneous group scan protocol (Altman 1974) was used to assess time spent in 1m proximity, friendly body contact, and grooming by all female-female dyads. Only one social activity with one partner was recorded per individual per scan. Priority was given to grooming over contact sitting and contact sitting over being in close proximity and if the subject was engaged in interaction with two partner in the same priority class, one partner was chosen randomly. These data were used to calculate relationship indices (see below). In 311 scans, 7746 individual behavioural scores were recorded, i.e. the average scan had almost 25 of the 29 individuals. Between scans, all-occurrence sampling (Altman 1974) was used to record agonistic interactions between adult females to establish the dominance hierarchy. In separate sampling sessions, we used sequence or event sampling (Altmann, 1974) to record the sequence of behaviours that unfolded around 1132 grooming interactions between two adult females where rank, kinship, and relationship information was available for both groomers. Opportunity to intervene in grooming was recorded for all females that were in the smaller indoor enclosure with the grooming dyad or all females that had an unobstructed view on the grooming dyad in the larger outdoor enclosure. Intervention was recorded for all behaviours that could affect the two grooming individuals including aggressive behaviours, affiliative behaviours, neutral approaches, and passing by at close range. The outcome of the intervention was recorded as effectively interrupting the grooming or not, and as yielding access to one of the grooming partners for active or passive grooming or contact sitting or none of these options. Observations ceased when all participants departed the grooming location.

### Variables

Dominance ranks were derived from normalized David’s Scores (De Vries et al., 2006) calculated from a winner-loser matrix of 1311 decided dyadic agonistic conflicts where one partner only showed submissive behaviour (bared-teeth display, crouch, give ground) either spontaneously or in reaction to aggression by the other partner, who did not show submission.

The strength of the affiliative relationship between two females was assayed with a dyadic composite sociality index (DSI, (Silk et al., 2013)) with three components: the number of scans including grooming (total 1564), contact sitting (total 1570), and close proximity (<1m; total 1412). The grooming included in the DSI was take from scans, and does therefore not contain the same bouts as those used to assess grooming interventions. All components were positively correlated in row-wise Kendall’s matrix correlations with 10,000 randomizations of the symmetric matrix columns in MatMan1.1 (Netto et al., 1993) at row-wise average tau of 0.38 to 0.44 and all p<0.001. The DSI by definition has an average of 1 and increases as dyads have affiliation times that are exceedingly the average dyad in the group.

Maternal kinship data were available for all females from the stud book of the colony. Dyads were classified as close kin if r≥0.25 and non-kin if r<0.25.

### Analyses

All models were fitted in R v4.0.0 (R Development Core Team & R Core Team, 2020) using the ‘brms’ (Bürkner, 2018) and ‘Rstan’ (Stan Development Team, 2020) packages. Posterior estimates were generated using the Hamiltonian Monte Carlo algorithm. We used 3000 iterations for two chains; chain convergence was assessed by visual inspection of traceplots (McElreath, 2018), showing no convergence problems. For all fixed effects in all models, we used weakly informative, t-distributed priors (Lemoine, 2019). All continuous variables were z-standardised (Schielzeth, 2010). The DSI relationship index was log-transformed in all models as it was highly skewed. Variance Inflation Factors were assessed to rule out collinearity problems (Field et al., 2012) using the R package ‘car’ (Fox et al., 2014). For all models, we present the 95% credible interval.

#### 1) Which grooming bouts faced interventions?

We tested whether certain characteristics make it more probable that a grooming bout sees an intervention. We fitted a Bayesian generalized linear multilevel model with a binomial response variable and logit link function. The response determined whether an intervention took place (n = 521) or not (n = 611). Data were structured to reflect the rank relations between the groomers. We included as random effects the identity of the higher-ranking individual and the lower-ranking individual, and defined variables by the same standard. Test predictors in this model were the rank of the higher-ranking individual and the rank of the lower-ranking individual. We also included whether the groomers were kin and their DSI. We included the number of bystanders as control variable. We included the log-transformed duration of the grooming bout as offset term. We included random intercepts for the two individuals; as most dyads had only one grooming bout, we did not include an intercept for the dyad. We included random slopes for the rank of the partner within each of the identities.

#### 2) Which bystander intervened?

For grooming bouts that showed interventions, we tested which bystander intervened. Thus, for 521 grooming bouts, we had overall 11,368 data points representing possible interveners (mean: 22 bystanders, range: 5 – 27). It is difficult representing the relationships between three individuals in a model in a meaningful way without influencing model fitting based on the specific solution selected. In a previous paper (Mielke et al., 2017), two models were run, one structured by the rank relations of the groomers, and one structured by their bond with the potential intervener; however, that way, the results for dominance rank and social relationships were separated. Here, we tested for the impact of rank, kinship, and affiliative relationship strength in one model, applying conditional logistic regression analysis (Kajokaite et al., 2019). If at least one individual necessarily intervened in each bout (as is the case in this model), then their rank, relationship to the groomers etc. matter not only globally, but also in relation to all other bystanders in that bout. In conditional logistic regression analysis, the importance of each effect on the outcome is dependent on the values for all other available choices in this bout, as well as their overall value.

We implemented conditional logistic regression analysis in a Bayesian framework using the ‘stan_logit’ function in the ‘rstanarm’ package (Goodrich et al., 2020). The outcome variable was binomial, whether a bystander intervened or not. Data were stratified by grooming bout identifier, so that all bystanders for a bout were tested against each other. As testing was conducted mainly within-bout, information about the two groomers did not vary and was not included in the model. For each bystander, we included as fixed effects their rank, the relationship value they had with each groomer, the absolute rank difference they had with each groomer (to test whether they disrupt grooming of close-ranked competitors), and whether they were close kin with each groomer. We included the random intercept for the bystander identity, with slopes for all main effects.

#### 3) Which groomer did interveners attempt to get access to?

For all grooming bouts for which a target could be identified (n = 411), we tested which of the two groomers was selected. The ‘target’ in this case is either the first individual the intervener grooms, or the groomer that is not supplanted or attacked by the intervener (Mielke et al., 2017). We again fitted a conditional logistic regression analysis, with each bout represented by the two groomers and their characteristics tested against each other. One groomer was always chosen (1), the other one not (0). Each data point was represented by the rank of the groomer (to test whether individuals select the higher-ranking groomer as target), the DSI between groomer and intervener, and whether they were kin related. We included a random intercept for the groomer identity, with slopes for the DSI and kinship.

#### 4) What determined intervention outcome?

Interventions can have different outcomes from the perspective of the intervener: no further grooming happens, grooming continues without them, or the intervener gains access to one or two grooming partners. In the absence of overt aggression in grooming interventions, as was the case here, the outcome is mainly determined not by the decision of the intervener, but by the willingness of the groomers to remain in close proximity to the intervener and participate in grooming with them. If neither groomer can tolerate the presence of the intervener, grooming ends; if one individual can tolerate the intervener’s presence, they groom her alone; if both are comfortable close to the intervener, polyadic grooming can occur. This level of tolerance for polyadic grooming was a major difference between chimpanzees and mangabeys (Mielke et al., 2017). Here, we tested the outcome of grooming interventions both from the perspective of the groomers and the intervener: when do groomers stay or leave? When does an intervener gain access to a grooming partner or disrupt a bout?

For all interventions where both groomers’ behaviour was identifiable (n = 504), Model 4.1 tests for each of the two groomers whether they remained in the grooming bout (1, n = 742) or not (0, n = 266), based on their relationship to the intervener. We assumed that the choices of the two groomers were independent from each other. In n = 102 bouts, both individuals left; in n = 62, one individual left; in n = 340 bouts, both individuals remained in the grooming bout post-intervention. We fitted a Generalized Linear Mixed Model with binomial error distribution. As fixed effects, we included the ranks of the groomer and the intervener and the interaction term between them, the kinship between the groomer and the intervener, and the DSI between the groomer and the intervener. As random effects, we included the intercepts ID of the groomer, the intervener, and the bout ID, and the slopes for the ranks and DSI within the individual IDs.

Model 4.2 tests for each grooming intervention which of the following outcomes were achieved: the intervener was ignored and grooming continued (n = 211), the bout was entirely disrupted (n = 102), or the intervener gained access to at least one grooming partner (n = 191). We fitted a multinomial logistic regression, again using the ‘brms’ and ‘rstan’ packages, using each of the three outcomes as possible response. As fixed effects, we included the intervener rank; a factor indicating whether the intervener was kin with either groomer; and a continuous variable indicating the maximum DSI they had with either groomer. As random effect, we included the identity of the intervener.

## Results

### Which grooming bouts faced interventions? (Model 1)

There was only weak impact of any test predictors on whether a grooming bout was subject to interventions (Tab. 1 for posterior distributions of fixed effects). There was weak evidence that with increasing rank of the higher-ranking and lower-ranking groomers, the probability of an intervention was reduced: the Odds Ratios showed that one standard deviation increase in the rank of the higher-ranking groomer reduced the odds by a factor of 0.81, and for the lower-ranking groomer by a factor of 0.82. There was little evidence that the relationship index or kinship of the groomers influenced intervention likelihood. For each standard deviation increase in number of bystanders, the likelihood of intervention increased by 18% (OR = 1.18). The full model explained a medium portion of variance in incidence of intervention R^2^ = 0.48 (Gelman et al. 2019).

**Table 1:** Results for Model 1, testing which grooming bouts received interventions. Posterior estimates and 95% credible interval for all fixed effects, and Odds Ratio for the estimates.

|  | Estimate | Estimate Error | Q2.5 | Q97.5 | Odds Ratio Estimate |
| --- | --- | --- | --- | --- | --- |
| Intercept | -0.35 | 0.12 | -0.58 | -0.12 | 0.71 |
| **Rank Higher-Ranking Groomer** | **-0.21** | **0.14** | **-0.49** | **0.06** | **0.81** |
| **Rank Lower-Ranking Groomer** | **-0.20** | **0.12** | **-0.44** | **0.04** | **0.82** |
| Relationship Strength Groomers | -0.08 | 0.11 | -0.29 | 0.13 | 0.93 |
| Kin Groomers | -0.11 | 0.27 | -0.64 | 0.41 | 0.89 |
| **Number of bystanders** | **0.16** | **0.08** | **0.02** | **0.31** | **1.18** |

### Which bystander intervened? (Model 2)

Several test predictors strongly influenced which of the bystanders intervened in grooming bouts (Table 2). The likelihood to intervene increased with bystander dominance rank, with a rank increase of one standard deviation increasing intervention likelihood by 78% (OR = 1.78, Fig. 1A). The rank difference between bystander and the higher-ranking groomer had no effect, but individuals were more likely to intervene when they were close in rank to the lower-ranking groomer (OR = 0.72, Fig. 1B). Individuals that were closely affiliated with either the higher-ranking (OR = 1.95, Fig. 1C) or lower-ranking groomer (OR= 1.61) had a higher probability of intervening, as did those that were kin related to either groomer (OR = 1.47 and OR = 1.89; Fig. 1D). The full model explained 8% of the variance in incidence of bystander intervention (R^2^ = 0.08).

**Table 2:** Results for Model 2, testing which bystanders intervened into grooming bouts. Posterior estimates and 95% credible interval for all fixed effects, and Odds Ratio for the estimates.

|  | Estimate | Estimate Error | Q2.5 | Q97.5 | Odds Ratio Estimate |
| --- | --- | --- | --- | --- | --- |
| **Rank Bystander** | **0.58** | **0.12** | **0.36** | **0.82** | **1.78** |
| Rank Difference to Higher-Ranking Groomer | 0.02 | 0.07 | -0.12 | 0.16 | 1.02 |
| **Rank Difference to Lower-Ranking Groomer** | **-0.33** | **0.11** | **-0.54** | **-0.12** | **0.72** |
| **Relationship Strength Bystander with Higher-Ranking Groomer** | **0.67** | **0.06** | **0.54** | **0.79** | **1.95** |
| **Relationship Strength Bystander with Lower-Ranking Groomer** | **0.47** | **0.06** | **0.36** | **0.58** | **1.61** |
| **Kinship Bystander with Higher-Ranking** | **0.39** | **0.15** | **0.08** | **0.69** | **1.47** |
| **Kinship Bystander with Lower-Ranking** | **0.64** | **0.18** | **0.29** | **0.99** | **1.89** |

**Figure 1:**
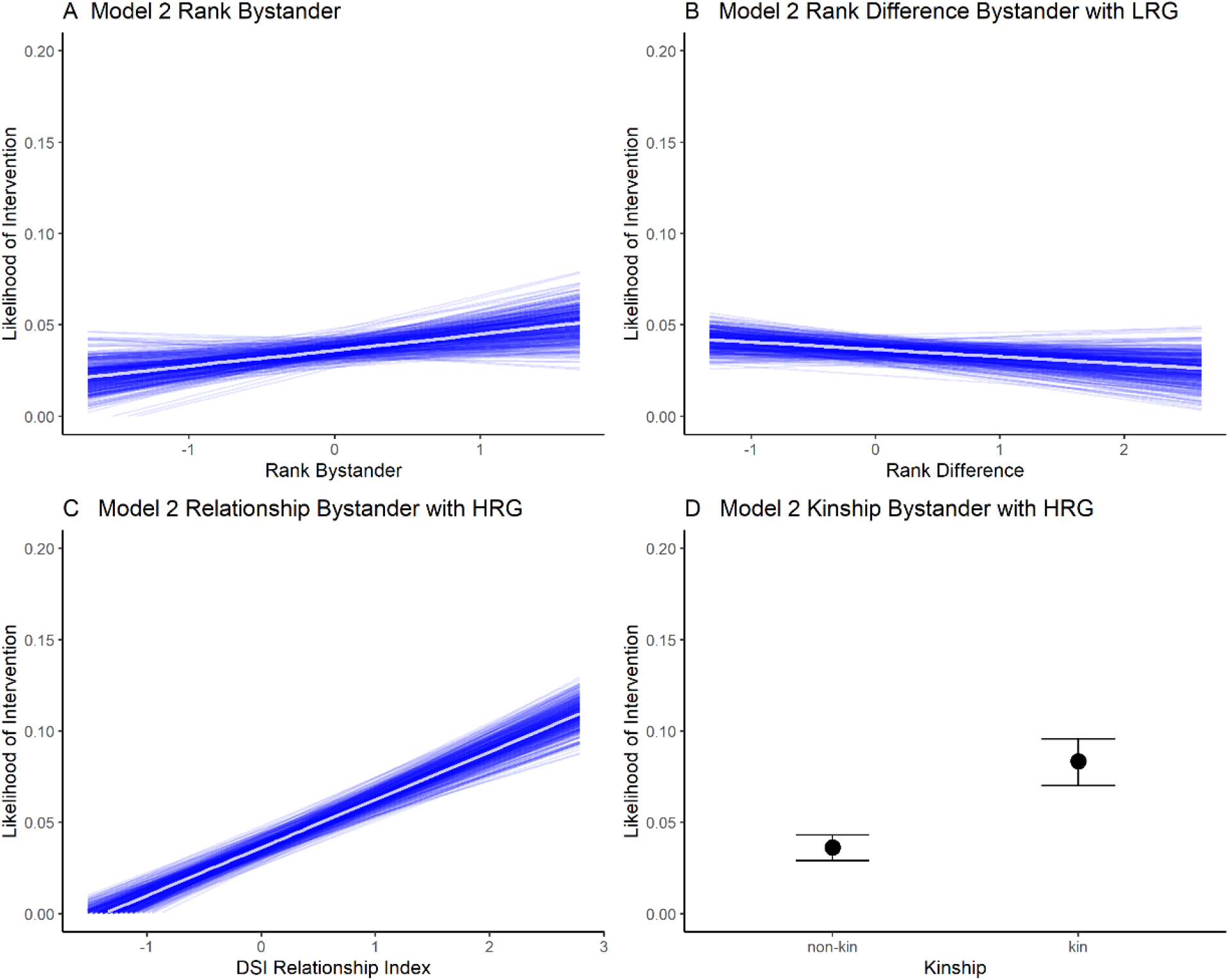
Impact of the bystander rank (A), rank difference between the bystander and the lower-ranking groomer (B), the relationship index of the bystander and the higher-ranking groomer (C) and kinship between bystander and higher-ranking groomer (D) on intervention likelihood (Model 2). Plots visualise impact of predictors for 400 model runs. Lines in spaghetti plots represent different draws of conditional effects of the model. Continuous predictors are z-standardised.

### Which groomer did interveners target? (Model 3)

Testing which of the two groomers the interveners sought to gain access to (Table 3), we found that they tended to target the groomer with whom they had a stronger affiliative relationships; one standard deviation increase in the relationship index increased the likelihood of choice by 43% (OR = 1.43, Fig. 6). There was no support for a bias towards higher-ranking groomers or kin. Together these predictors explained 13% of variance in the response (R^2^ = 0.13).

**Table 3:** Results for Model 3, testing which groomer was targeted by intervention. Posterior estimates and 95% credible interval for all fixed effects, and Odds Ratio for the estimates.

|  | Estimate | Estimate Error | Q2.5 | Q97.5 | Odds Ratio Estimate |
| --- | --- | --- | --- | --- | --- |
| Rank Groomer | 0.07 | 0.11 | -0.13 | 0.30 | 1.08 |
| **Relationship Strength Groomer – Intervener** | **0.30** | **0.12** | **0.08** | **0.54** | **1.35** |
| Kin Groomer – Intervener | 0.11 | 0.25 | -0.40 | 0.58 | 1.12 |

### What determined intervention outcome? (Model 4)

We tested what determined the outcome of the intervention, i.e. whether groomers avoided the intervener and left the scene or remained to continue the grooming bout (Model 4.1, Tab. 4). Decreasing the interveners rank strongly increased the likelihood that groomers stayed (OR = 0.65, Fig. 2A), especially with increasing groomer rank (OR = 1.17), while the interaction between the two terms did not strongly affect staying likelihood. With increasing affiliative relationship strength between the groomer and the intervener, groomers were more likely to stay (OR = 1.32, Fig. 2B). There was no evidence that kin relations to the intervener informed groomers’ decision to leave or stay. The effect size of this model was R^2^ = 0.08.

**Table 4:**
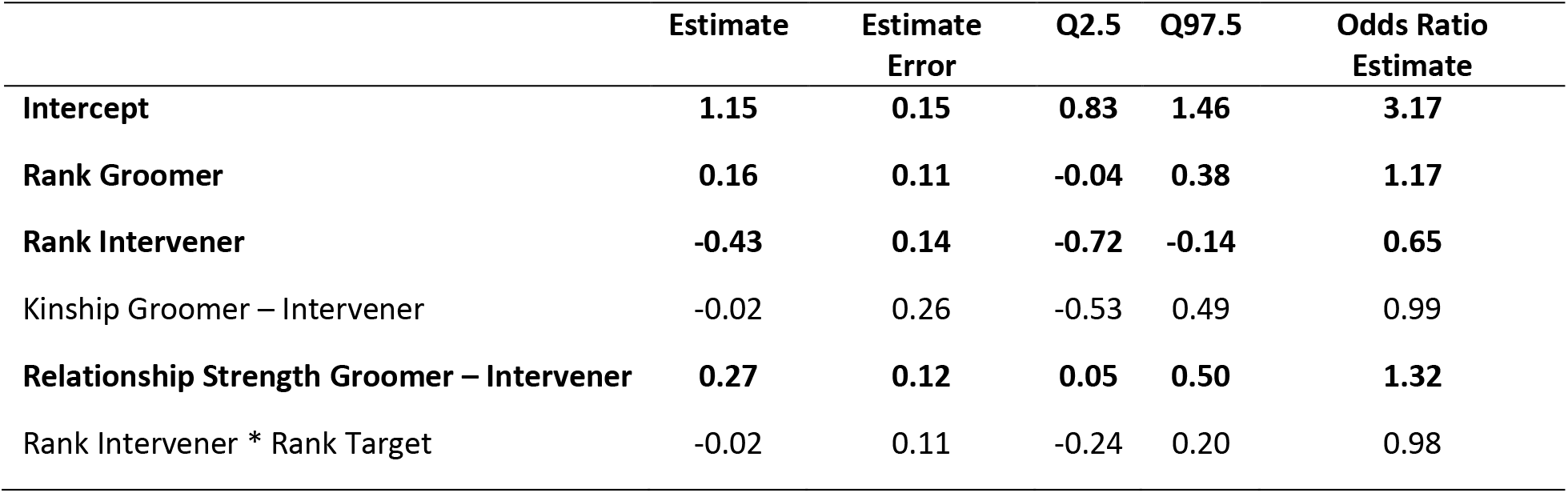
Results for Model 4.1, testing which groomers remained in the grooming bout after the intervention. Posterior estimates and 95% credible interval for all fixed effects, and Odds Ratio for the estimates.

**Figure 2:**
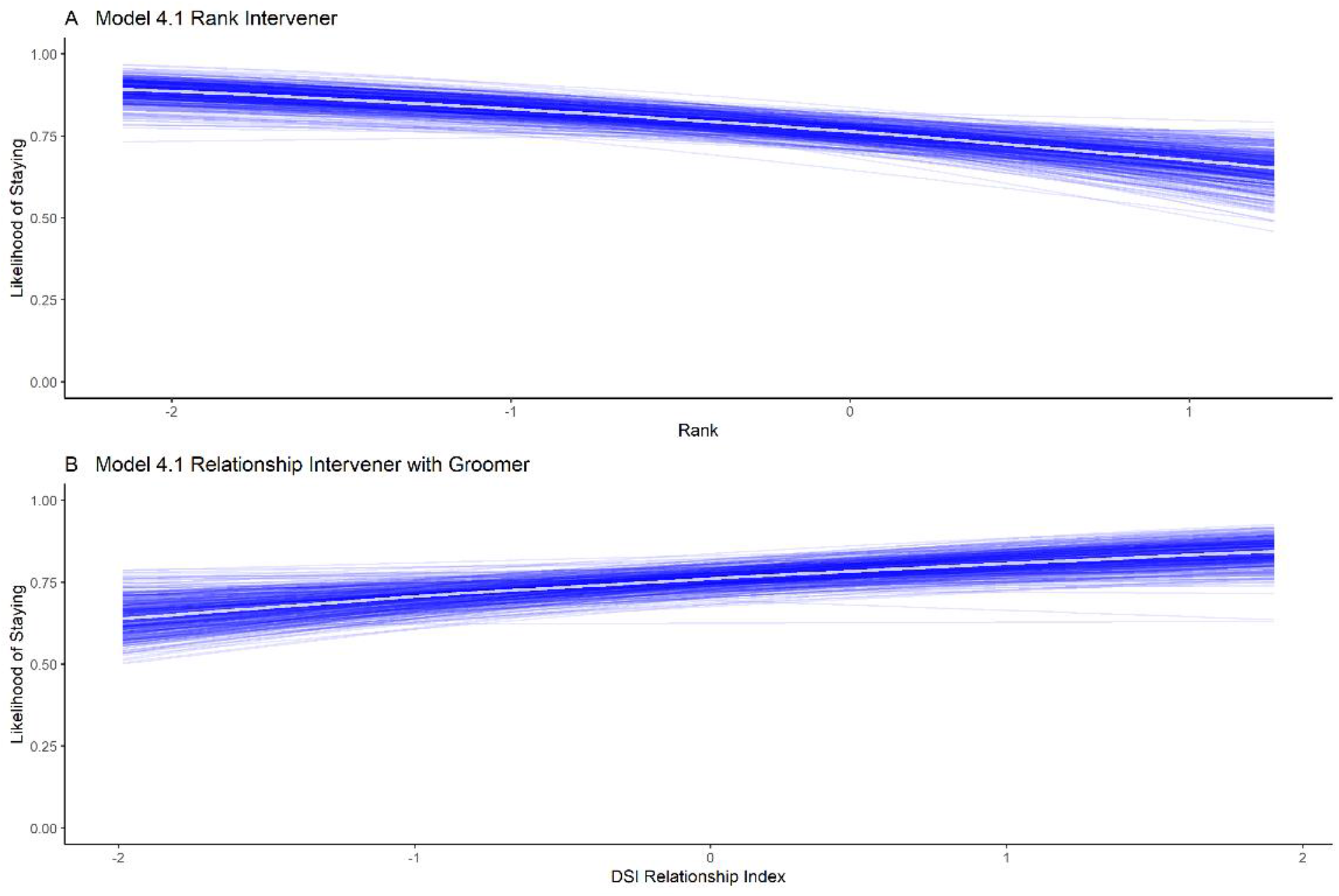
Impact of intervener rank (A) and the relationship index between intervener and groomer (B) on whether groomers stayed part of the grooming bout after an intervention (Model 4.1). Spaghetti plots of 400 model runs. Lines represent different draws of conditional effects of the model. Predictors were z-standardised.

For Model 4.2, we reversed the question and tested what determine whether the intervener was ignored by the groomers, disrupted the grooming bout, or gained access to at least one grooming partner (Tab. 5). There was consistent evidence that with increasing affiliative relationship strength with at least one groomer, interveners were more likely to gain access to at least one grooming partner (OR = 1.50) rather than be ignored, and they were less likely to disrupt the bout than be ignored (OR = 0.70, Fig. 3A). An increase of one standard deviation of intervener rank increased the likelihood to disrupt the bout rather than be ignored by 40% (OR = 1.40, Fig. 3B). These results mirror those of Model 4.1.

**Table 5:** Posterior estimates and 95% credible interval for all fixed effects of Model 4.2, testing whether grooming interventions resulted in disruption, access to a groomer, or being ignored. The reference category in this case is ‘being ignored’.

|  | Estimate | Estimate Error | Q2.5 | Q97.5 | Odds Ratio Estimate |
| --- | --- | --- | --- | --- | --- |
| Intercept - Access | 0.04 | 0.24 | -0.43 | 0.52 | 1.04 |
| Intercept - Disruption | -0.85 | 0.23 | -1.29 | -0.40 | 0.43 |
| Access – Kinship Intervener with either Groomer | -0.36 | 0.28 | -0.91 | 0.18 | 0.70 |
| Access – Rank Intervener | 0.08 | 0.18 | -0.26 | 0.44 | 1.08 |
| **Access – Maximum Relationship Strength with either Groomer** | **0.41** | **0.16** | **0.11** | **0.72** | **1.50** |
| Disruption – Kinship Intervener with either Groomer | -0.07 | 0.31 | -0.68 | 0.53 | 0.93 |
| **Disruption – Rank Intervener** | **0.34** | **0.16** | **0.03** | **0.68** | **1.41** |
| **Disruption – Maximum Relationship Strength with either Groomer** | **-0.36** | **0.16** | **-0.67** | **0.06** | **0.70** |

**Figure 3:**
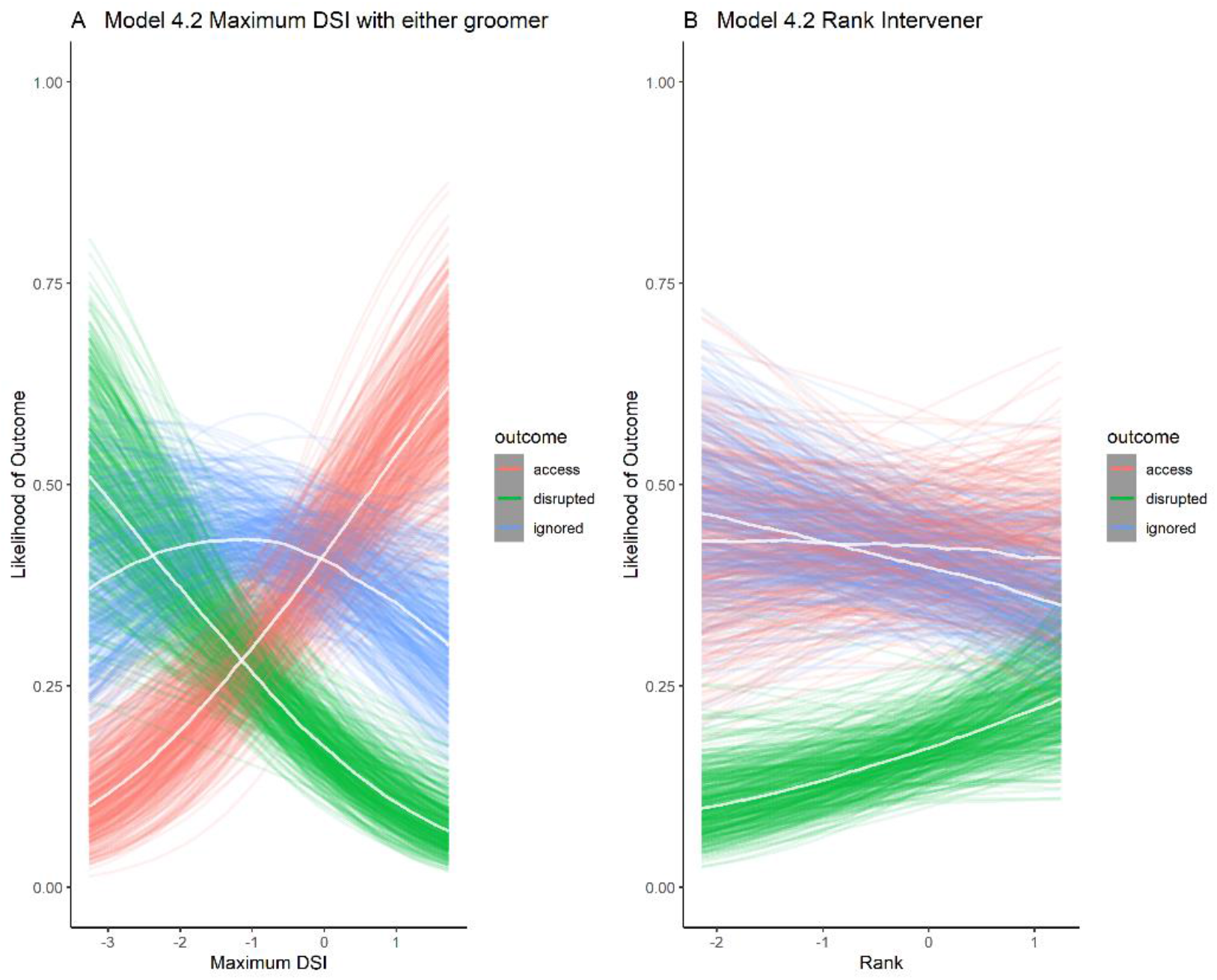
Impact of the maximum relationship index of the intervener with either groomer (A) and intervener rank (B) on the outcome of the intervention (Model 4.2). Spaghetti plots of 400 model runs. Lines represent different draws of conditional effects of the model; white lines within the distribution represent the median of all draws. Predictors were z-standardised.

## Discussion

Being able to manipulate the social interactions and potential relationships of other group members gives animals considerable influence over their own fate in groups of interdependent agents. This is a process of social niche construction: Individuals modify the selective context – here, the social interactions of all individuals in the group – in which they have to make adaptive decisions (Ryan et al., 2016). While we know more and more about the ability of animal to monitor relationships between others (Berthier & Semple, 2018; Schülke et al., 2020), the question remains how flexible they are in employing this knowledge to their own immediate gain and to manipulate interactions between others. In an aggressive context, policing (Beisner & McCowan, 2013; Flack et al., 2006), agonistic intervention (Barrett et al., 2015), and third-party reconciliation (de Waal & van Roosmalen, 1979; Wittig & Boesch, 2005) have been described as ways for individuals to influence group members. There is increasing evidence that similar strategies exist for affiliative behaviour (Massen et al., 2014; Mondragón-Ceballos, 2001; Schneider & Krueger, 2012; Ward et al., 2009). In this study, we examined how multiple individual and relationship characteristics (rank, kinship, and affiliative relationships) influence decisions about interventions in female rhesus macaque grooming.

Through a series of models, we explored which grooming dyads faced interventions, by whom, which groomer the intervener tried to gain access to, and what determined intervention outcomes. To understand the wealth of results, it pays to first focus on the main difference previously found between sooty mangabeys and chimpanzees (Mielke et al., 2017): In mangabeys, grooming is never triadic, so all interventions were disruptive, which favoured high-ranking individuals and allowed them to manipulate grooming bouts of close affiliates and closely-ranked competitors. This seems to also be the case in stumptailed macaques (Mondragón-Ceballos, 2001) and mandrills (Schino & Lasio, 2018). In chimpanzees, almost all interveners joined the grooming bout, as polyadic grooming is common (Nakamura, 2003), making intervention success independent of rank. Who decided to intervene, however, did not seem to differ between species; what differed was the ability of multiple individuals to be engaged in the same interaction. Rhesus macaques in our study group showed both joining and disruptive interventions, allowing us to test what determined the outcome of grooming interventions.

We found broad similarities between rhesus macaques and previous results on affiliation interventions in other species. As in mangabeys and mandrills (Mielke et al., 2017; Schino & Lasio, 2018), high-ranking bystanders were more likely to intervene, as they were less likely to be ignored than low-ranking interveners. Consequently, high-ranking grooming dyads saw slightly fewer interventions overall. Rank therefore seemed to confer power to individuals in deciding when to groom and intervene (Barrett et al., 1999; Sambrook et al., 1995). As in mangabeys and chimpanzees, bystanders were more likely to intervene when the lower-ranking groomer was close to them in rank, potentially to disrupt bond-formation of closely ranked competitors (Mielke et al., 2017). Bystanders were more likely to intervene when they had strong affiliative relationships with either groomer, as in horses (Schneider & Krueger, 2012), mangabeys, and chimpanzees. Kinship with either groomer also played a strong role. Like mangabeys and chimpanzees, our subjects did not seem to target the higher-ranking groomer as subsequent grooming partner (Mielke et al., 2017). The consistent finding across species of individuals influencing affiliative interactions of their own bond partners raises further questions as to how interventions relate to the evolution of jealousy (Harris & Prouvost, 2014). To establish the presence of friendship or romantic jealousy in humans observationally, we would test how we prevent partners from establishing new connections, and renewed commitment with the partner (Aune & Comstock, 1991). Losing a close affiliate affects the life of animals with long-lasting bonds, and having a ‘warning system’ to detect and the behavioural means to prevent such loss might prove valuable. Future studies of the detailed reaction of bystanders to observed cooperation are necessary to determine whether outward signs of agitation are detectable in primates (Harris & Prouvost, 2014).

We found that rhesus macaques used interventions to gain access to close affiliates, in contrast to mangabeys, chimpanzees, and domestic dogs (Ward et al., 2009). Strong affiliative relationships with either groomer increased the likelihood of grooming access post-intervention, as groomers were less likely to leave. Thus, rhesus macaque females stop affiliates from grooming others and get access to them for additional grooming. These two functions are not mutually exclusive, and targeting affiliates could indeed function to repair potential damage to the relationship. Alternatively, low-ranking individuals might use their affiliative and kin relationships to intervene in grooming bouts and access partners that they would not be able to, based on rank alone. An indication for this is that individuals were more likely to intervene if they were genetically related to either groomer. At the same time, kin were not more likely to be targeted by interventions or to remain part of the grooming bout post-intervention. Interveners were more likely to gain access to grooming bouts if they had an affiliative relationship with at least one groomer and many of the subsequent grooming bouts were polyadic. Interventions might therefore function to increase an individual’s grooming network, by using close affiliates and kin to access other group members; again shaping individuals’ own social niche.

These results shine another light on the importance of affiliation between more than two partners. In many primate species, dyadic grooming seems to be the norm or only option. However, this artificially limits access to grooming partners: if individuals were able to groom with more partners at the same time, many of the time restrictions that are thought to underlie competition for grooming partners (Sambrook et al., 1995) would disappear. Some species, such as chimpanzees and bonobos, regularly groom as clusters (Nakamura, 2003; Sakamaki, 2013), which could allow individuals to extend their social network compared to species with strict priority of access; this seemed to be the case in the rhesus macaques. The question is what underlies this eschewal of polyadic grooming: whether this is a question of multiple individuals tolerating each other in close distance (Jaeggi et al., 2010), whether coordinating more than two individuals poses a cognitive challenge (Wacewicz et al., 2017), or whether keeping track of reciprocal exchanges becomes more difficult with increasing groomer numbers (Schino & Aureli, 2009). Here, individuals were more likely to remain and continue grooming if they were affiliated to the intervener or the intervener was low in rank; both conditions that reduce the likelihood of aggression. Thus, tolerance between individuals might be an important aspect underlying polyadic grooming, rather than potential cognitive difference between species.

Theories regarding the evolution of cognitive abilities in animals often focus on the challenges posed by an ever-changing social environment and the need to track and adapt to the actions of other agents (Seyfarth & Cheney, 2015). Social bonds (Ellis et al 2019) and indirect connections to third parties (Brent, 2015) are highly relevant for rhesus macaques. Grooming is common and an important tool to navigate relationships (Balasubramaniam & Berman, 2017). They track who is close to them (Semple et al., 2009) and what these individuals are doing (Schülke et al., 2020). Here, we show that rather than simply gathering information, rhesus macaques use their own behaviour to manipulate interactions between others to their own benefits and construct their own social niche. Females based their decisions during interventions on a number of characteristics of the groomers, their relationship to these individuals, but also potentially triadic awareness of the relationship between the groomers (Kubenova et al., 2016; Wittig et al., 2014). Thus, this study presents additional evidence for the flexibility and cognitive skills primates use when navigating and shaping their social environment.

## Data Availability

Data and scripts: https://github.com/AlexMielke1988/Mielke-et-al-Rhesus-Interventions

## Acknowledgements

We thank Uwe Schönmann, Annette Husung, and the animal keepers at the German Primate Center for logistical support. The study benefitted from discussions in the DFG-RTG 2070 Understanding Social Relationships (DFG, German Research Foundation Project number 254142454). AM was funded by a British Academy Newton International Fellowship. We thank Kotrina Kajokaite for helpful discussions of conditional logistic regression models.

## Notes

https://github.com/AlexMielke1988/Mielke-et-al-Rhesus-Interventions

### Competing Interest Statement

The authors have declared no competing interest.

## References

Altmann, J. (1974). Observational Study of Behavior: Sampling Methods. Behaviour, 49(3–4), 227–266. https://doi.org/10.1163/156853974X00534

Archie, E. A., Tung, J., Clark, M., Altmann, J., & Alberts, S. C. (2014). Social affiliation matters: both same-sex and opposite-sex relationships predict survival in wild female baboons. Proceedings of the Royal Society of London B: Biological Sciences, 281(1793), 20141261. https://doi.org/10.1098/rspb.2014.1261

Aune, K. S., & Comstock, J. (1991). Experience and expression of jealousy: comparison between friends and romantics. Psychological Reports, 69(1), 315–319. https://doi.org/10.2466/pr0.1991.69.1.315

Balasubramaniam, K. N., & Berman, C. M. (2017). Grooming interchange for resource tolerance: Biological markets principles within a group of free-ranging rhesus macaques. Behaviour, 154(11), 1145–1176. https://doi.org/10.1163/1568539X-00003462

Barrett, L., Flinn, M., De Waal, F. B. M., Perry, S., Mitani, J. C., Gavrilets, S., & Bissonnette, A. (2015). Coalitions in theory and reality: A review of pertinent variables and processes. In Behaviour (Vol. 152, Issue 1, pp. 1–56). https://doi.org/10.1163/1568539X-00003241

Barrett, L., Henzi, S. P., Weingrill, T., Lycett, J. E., & Hill, R. A. (1999). Market forces predict grooming reciprocity in female baboons. Proceedings of the Royal Society B: Biological Sciences, 266(1420), 665. https://doi.org/10.1098/rspb.1999.0687

Barrett, L., Peter Henzi, S., & Lusseau, D. (2012). Taking sociality seriously: The structure of multi-dimensional social networks as a source of information for individuals. Philosophical Transactions of the Royal Society B: Biological Sciences, 367(1599), 2108–2118. https://doi.org/10.1098/rstb.2012.0113

Beisner, B. A., & McCowan, B. (2013). Policing in Nonhuman Primates: Partial Interventions Serve a Prosocial Conflict Management Function in Rhesus Macaques. PLoS ONE, 8(10), e77369. https://doi.org/10.1371/journal.pone.0077369

Berthier, J. M., & Semple, S. (2018). Observing grooming promotes affiliation in Barbary macaques. Proceedings of the Royal Society B: Biological Sciences, 285(1893), 20181964. https://doi.org/10.1098/rspb.2018.1964

Brent, L. J. N. (2015). Friends of friends: Are indirect connections in social networks important to animal behaviour? Animal Behaviour, 103, 211–222. https://doi.org/10.1016/j.anbehav.2015.01.020

Bürkner, P. C. (2018). Advanced Bayesian multilevel modeling with the R package brms. R Journal, 10(1), 395–411. https://doi.org/10.32614/rj-2018-017

Cameron, E. Z., Setsaas, T. H., & Linklater, W. L. (2009). Social bonds between unrelated females increase reproductive success in feral horses. Proceedings of the National Academy of Sciences of the United States of America, 106(33), 13850–13853. https://doi.org/10.1073/pnas.0900639106

Carter, G. G., Farine, D. R., Crisp, R. J., Vrtilek, J. K., Ripperger, S. P., & Page, R. A. (2020). Development of New Food-Sharing Relationships in Vampire Bats. Current Biology, 30(7), 1275–1279.e3. https://doi.org/10.1016/j.cub.2020.01.055

Carter, G. G., & Wilkinson, G. S. (2013). Food sharing in vampire bats: reciprocal help predicts donations more than relatedness or harassment. Proceedings of the Royal Society B: Biological Sciences, 280(1753), 20122573. https://doi.org/10.1098/rspb.2012.2573

Cords, M., & Aureli, F. (2000). Reconciliation and relationship quality. In F. B. M. de Waal & F. Aureli (Eds.), Natural conflict resolution (pp. 177–198). University of California Press.

De Moor, D., Roos, C., Ostner, J., & Schülke, O. (2020). Female Assamese macaques bias their affiliation to paternal and maternal kin. Behavioral Ecology, 31(2), 493–507. https://doi.org/10.1093/beheco/arz213

De Vries, H., Stevens, J. M. G., & Vervaecke, H. (2006). Measuring and testing the steepness of dominance hierarchies. Animal Behaviour, 71(3), 585–592. https://doi.org/10.1016/j.anbehav.2005.05.015

de Waal, F. B. M., & van Roosmalen, A. (1979). reconciliation and consolation among chimpanzees. Behavioral Ecology and Sociobiology. https://doi.org/10.1007/BF00302695

Ellis, S., Snyder-Mackler, N., Ruiz-Lambides, A., Platt, M. L., & Brent, L. J. N. (2019). Deconstructing sociality: the types of social connections that predict longevity in a group-living primate. Proceedings of the Royal Society B: Biological Sciences, 286(1917), 20191991. https://doi.org/10.1098/rspb.2019.1991

Field, A. P., Miles, J., & Field, Z. (2012). Discovering statistics using R. In Choice Reviews Online (Vol. 50, Issue 04). Sage publications. https://doi.org/10.1111/insr.12011_21

Flack, J. C., Girvan, M., De Waal, F. B. M., & Krakauer, D. C. (2006). Policing stabilizes construction of social niches in primates. Nature, 439(7075), 426–429. https://doi.org/10.1038/nature04326

Fox, J., Weisberg, S., Adler, D., Bates, D. M., Baud-Bovy, G., Ellison, S., Firth, D., Friendly, M., Gorjanc, G., Graves, S., Heiberger, R., Laboissiere, R., Mon, G., Murdoch, D., Nilsson, H., Ogle, D., Ripley, B., Venables, W., Zeileis, A., & R Development Core Team. (2014). Package ‘car.’ R Topics Documented, 167.

Goodrich, B., Gabry, J., Ali, I., & Brilleman, S. (2020). rstanarm: Bayesian applied regression modeling via Stan.

Harris, C. R., & Prouvost, C. (2014). Jealousy in dogs. PLoS ONE, 9(7), e94597. https://doi.org/10.1371/journal.pone.0094597

Jaeggi, A. V., Stevens, J. M. G., & Van Schaik, C. P. (2010). Tolerant food sharing and reciprocity is precluded by despotism among bonobos but not chimpanzees. American Journal of Physical Anthropology, 143(1), 41–51. https://doi.org/10.1002/ajpa.21288

Jolly, A. (1966). Lemur social behavior and primate intelligence. Science, 153(3735), 501–506. https://doi.org/10.1126/science.153.3735.501

Kajokaite, K., Whalen, A., Panchanathan, K., & Perry, S. (2019). White-faced capuchin monkeys use both rank and relationship quality to recruit allies. Animal Behaviour, 154, 161–169. https://doi.org/10.1016/j.anbehav.2019.06.008

Kalbitz, J., Ostner, J., & Schülke, O. (2016). Strong, equitable and long-term social bonds in the dispersing sex in Assamese macaques. Animal Behaviour, 113, 13–22. https://doi.org/10.1016/j.anbehav.2015.11.005

Kubenova, B., Konecna, M., Majolo, B., Smilauer, P., Ostner, J., & Schülke, O. (2016). Triadic awareness predicts partner choice in male-infant-male interactions in Barbary macaques. Animal Cognition, 20(2), 1–12. https://doi.org/10.1007/s10071-016-1041-y

Lemoine, N. P. (2019). Moving beyond noninformative priors: why and how to choose weakly informative priors in Bayesian analyses. Oikos, 128(7), 912–928. https://doi.org/10.1111/oik.05985

Massen, J. J. M., & Sterck, E. H. M. (2013). Stability and Durability of Intra- and Intersex Social Bonds of Captive Rhesus Macaques (Macaca mulatta). International Journal of Primatology, 34(4), 770–791. https://doi.org/10.1007/s10764-013-9695-7

Massen, J. J. M., Szipl, G., Spreafico, M., & Bugnyar, T. (2014). Ravens intervene in others’ bonding attempts. Current Biology, 24(22), 2733–2736. https://doi.org/10.1016/j.cub.2014.09.073

McElreath, R. (2018). Statistical rethinking: A bayesian course with examples in R and stan. In Statistical Rethinking: A Bayesian Course with Examples in R and Stan. Chapman & Hall/CRC. https://doi.org/10.1201/9781315372495

Mielke, A., Preis, A., Samuni, L., Gogarten, J. F., Wittig, R. M., & Crockford, C. (2018). Flexible decision-making in grooming partner choice in sooty mangabeys and chimpanzees. Royal Society Open Science, 5(7), 172143. https://doi.org/10.1098/rsos.172143

Mielke, A., Samuni, L., Preis, A., Gogarten, J. F., Crockford, C., & Wittig, R. M. (2017). Bystanders intervene to impede grooming in western chimpanzees and sooty mangabeys. Royal Society Open Science, 4(11), 171296. https://doi.org/10.1098/rsos.171296

Mondragón-Ceballos, R. (2001). Interfering in affiliations: sabotaging by stumptailed macaques, Macaca arctoides. Animal Behaviour, 62(DECEMBER 2001), 1179–1187. https://doi.org/10.1006/anbe.2001.1861

Nakamura, M. (2003). “Gathering” of social grooming among wild chimpanzees: Implications for evolution of sociality. Journal of Human Evolution, 44(1), 59–71. https://doi.org/10.1016/S0047-2484(02)00194-X

Netto, W. J., Hanegraaf, P. L. H., & DE VRIES, H. (1993). Matman: a Program for the Analysis of Sociometric Matrices and Behavioural Transition Matrices. Behaviour, 125(3–4), 157–175. https://doi.org/10.1163/156853993x00218

Newton-Fisher, N. E., & Kaburu, S. S. K. (2017). Grooming decisions under structural despotism: the impact of social rank and bystanders among wild male chimpanzees. Animal Behaviour, 128, 153–164. https://doi.org/10.1016/j.anbehav.2017.04.012

Noë, R., & Hammerstein, P. (1995). Biological markets. Trends in Ecology & Evolution, 10(8), 336–339. https://doi.org/10.1016/S0169-5347(00)89123-5

Ostner, J., & Schülke, O. (2018). Linking Sociality to Fitness in Primates: A Call for Mechanisms. In Advances in the Study of Behavior (Vol. 50, pp. 127–175). Academic Press Inc. https://doi.org/10.1016/bs.asb.2017.12.001

Preis, A., Samuni, L., Mielke, A., Deschner, T., Crockford, C., & Wittig, R. M. (2018). Urinary oxytocin levels in relation to post-conflict affiliations in wild male chimpanzees (Pan troglodytes verus). Hormones and Behavior. https://doi.org/10.1016/j.yhbeh.2018.07.009

R Development Core Team, & R Core Team. (2020). R: A language and environment for statistical computing. R Foundation for Statistical Computing Vienna Austria, 0, {ISBN} 3-900051-07-0. https://doi.org/10.1038/sj.hdy.6800737

Ryan, P. A., Powers, S. T., & Watson, R. A. (2016). Social niche construction and evolutionary transitions in individuality. Biology and Philosophy, 31(1), 59–79. https://doi.org/10.1007/s10539-015-9505-z

Sakamaki, T. (2013). Social grooming among wild bonobos (Pan paniscus) at Wamba in the Luo Scientific Reserve, DR Congo, with special reference to the formation of grooming gatherings. Primates, 54(4), 349–359. https://doi.org/10.1007/s10329-013-0354-6

Sambrook, T. D., Whiten, A., & Strum, S. C. (1995). Priority of access and grooming patterns of females in a large and a small group of olive baboons. Animal Behaviour, 50(6), 1667–1682. https://doi.org/10.1016/0003-3472(95)80020-4

Samuni, L., Preis, A., Mielke, A., Deschner, T., Wittig, R. M., & Crockford, C. (2018). Social bonds facilitate cooperative resource sharing in wild chimpanzees. Proceedings of the Royal Society B: Biological Sciences, 285(1888), 20181643. https://doi.org/10.1098/rspb.2018.1643

Schielzeth, H. (2010). Simple means to improve the interpretability of regression coefficients. Methods in Ecology and Evolution, 1(2), 103–113. https://doi.org/10.1111/j.2041-210x.2010.00012.x

Schino, G., & Aureli, F. (2009). Chapter 2 Reciprocal Altruism in Primates. Partner Choice, Cognition, and Emotions. Advances in the Study of Behavior, 39(09), 45–69. https://doi.org/10.1016/S0065-3454(09)39002-6

Schino, G., & Lasio, F. (2018). Competition for grooming partners and interference in affiliation among female mandrills. Ethology, 124(8), 600–608. https://doi.org/10.1111/eth.12763

Schneider, G., & Krueger, K. (2012). Third-party interventions keep social partners from exchanging affiliative interactionswith others. Animal Behaviour, 83(2), 377–387. https://doi.org/10.1016/j.anbehav.2011.11.007

Schülke, O., Bhagavatula, J., Vigilant, L., & Ostner, J. (2010). Social bonds enhance reproductive success in male macaques. Current Biology, 20(24), 2207–2210. https://doi.org/10.1016/j.cub.2010.10.058

Schülke, O., Dumdey, N., & Ostner, J. (2020). Selective attention for affiliative and agonistic interactions of dominants and close affiliates in macaques. Scientific Reports, 10(1), 5962. https://doi.org/10.1038/s41598-020-62772-8

Semple, S., Gerald, M. S., & Suggs, D. N. (2009). Bystanders affect the outcome of mother-infant interactions in rhesus macaques. Proceedings. Biological Sciences / The Royal Society, 276(1665), 2257–2262. https://doi.org/10.1098/rspb.2009.0103

Seyfarth, R. M., & Cheney, D. L. (2015). Social cognition. Animal Behaviour, 103, 191–202. https://doi.org/10.1016/j.anbehav.2015.01.030

Silk, J. B., Alberts, S. C., Altmann, J., Cheney, D. L., & Seyfarth, R. M. (2012). Stability of partner choice among female baboons. Animal Behaviour, 83(6), 1511–1518. https://doi.org/10.1016/j.anbehav.2012.03.028

Silk, J. B., Beehner, J. C., Bergman, T. J., Crockford, C., Engh, A. L., Moscovice, L. R., Wittig, R. M., Seyfarth, R. M., & Cheney, D. L. (2009). The benefits of social capital: close social bonds among female baboons enhance offspring survival. Proceedings of the Royal Society B: Biological Sciences, 276(June), 3099–3104. https://doi.org/10.1098/rspb.2009.0681

Silk, J. B., Beehner, J. C., Bergman, T. J., Crockford, C., Engh, A. L., Moscovice, L. R., Wittig, R. M., Seyfarth, R. M., & Cheney, D. L. (2010). Female chacma baboons form strong, equitable, and enduring social bonds. Behavioral Ecology and Sociobiology, 64(11), 1733–1747. https://doi.org/10.1007/s00265-010-0986-0

Silk, J. B., Cheney, D., & Seyfarth, R. (2013). A practical guide to the study of social relationships. Evolutionary Anthropology, 22(5), 213–225. https://doi.org/10.1002/evan.21367

Snyder-Mackler, N., Burger, J. R., Gaydosh, L., Belsky, D. W., Noppert, G. A., Campos, F. A., Bartolomucci, A., Yang, Y. C., Aiello, A. E., O’Rand, A., Harris, K. M., Shively, C. A., Alberts, S. C., & Tung, J. (2020). Social determinants of health and survival in humans and other animals. In Science (New York, N.Y.) (Vol. 368, Issue 6493). NLM (Medline). https://doi.org/10.1126/science.aax9553

Städele, V., Roberts, E. R., Barrett, B. J., Strum, S. C., Vigilant, L., & Silk, J. B. (2019). Male–female relationships in olive baboons (Papio anubis): Parenting or mating effort? Journal of Human Evolution, 127, 81–92. https://doi.org/10.1016/j.jhevol.2018.09.003

Stan Development Team. (2020). RStan: the R interface to Stan.

Thierry, B., Aureli, F., Nunn, C. L., Petit, O., Abegg, C., & de Waal, F. B. M. (2008). A comparative study of conflict resolution in macaques: insights into the nature of trait covariation. Animal Behaviour, 75(3), 847–860. https://doi.org/10.1016/j.anbehav.2007.07.006

VanDierendonck, M. C., de Vries, H., Schilder, M. B. H., Colenbrander, B., Thorhallsdóttir, A. G., & Sigurjónsdóttir, H. (2009). Interventions in social behaviour in a herd of mares and geldings. Applied Animal Behaviour Science, 116(1), 67–73. https://doi.org/10.1016/j.applanim.2008.07.003

Wacewicz, S., Żywiczyński, P., & Chiera, A. (2017). An evolutionary approach to low-level conversational cooperation. Language Sciences, 63, 91–104. https://doi.org/10.1016/j.langsci.2017.01.005

Ward, C., Trisko, R. K., & Smuts, B. B. (2009). Third-party interventions in dyadic play between littermates of domestic dogs, Canis lupus familiaris. Animal Behaviour, 78(5), 1153–1160. https://doi.org/10.1016/j.anbehav.2009.07.033

Wittig, R. M., & Boesch, C. (2005). How to repair relationships - Reconciliation in wild chimpanzees (Pan troglodytes). Ethology, 111(8), 736–763. https://doi.org/10.1111/j.1439-0310.2005.01093.x

Wittig, R. M., Crockford, C., Langergraber, K. E., & Zuberbühler, K. (2014). Triadic social interactions operate across time: a field experiment with wild chimpanzees. Proceedings of the Royal Society B: Biological Sciences, 281(1779), 20133155. https://doi.org/10.1098/rspb.2013.3155

Young, C., Majolo, B., Schülke, O., & Ostner, J. (2014). Male social bonds and rank predict supporter selection in cooperative aggression in wild Barbary macaques. Animal Behaviour, 95, 23–32. https://doi.org/10.1016/j.anbehav.2014.06.007

